# Reference-free comparison of microbial communities via de Bruijn graphs

**DOI:** 10.1101/055020

**Authors:** Serghei Mangul, David Koslicki

## Abstract

Microbial communities inhabiting the human body exhibit significant variability across different individuals and tissues, and are suggested to play an important role in health and disease. High-throughput sequencing offers unprecedented possibilities to profile microbial community composition, but limitations of existing taxonomic classification methods (including incompleteness of existing microbial reference databases) limits the ability to accurately compare microbial communities across different samples. In this paper, we present a method able to overcome these limitations by circumventing the classification step and directly using the sequencing data to compare microbial communities. The proposed method provides a powerful reference-free way to assess differences in microbial abundances across samples. This method, called EMDeBruijn, condenses the sequencing data into a de Bruijn graph. The Earth Mover's Distance (EMD) is then used to measure similarities and differences of the microbial communities associated with the individual graphs. We apply this method to RNA-Seq data sets from a coronary artery calcification (CAC) study and shown that EMDeBruijn is able to differentiate between case and control CAC samples while utilizing all the candidate microbial reads. We compare these results to current reference-based methods, which are shown to have a limited capacity to discriminate between case and control samples. We conclude that this reference-free approach is a viable choice in comparative metatranscriptomic studies.

## 1. INTRODUCTION

The human microbiome, sometimes [17] referred to as “the forgotten organ,” contains a significantly larger number of genes than predicted in the human genome [6] and an increasing number of studies investigate its role in health and disease [9]. Traditional cultured-based approaches to microbial community profiling are limited in their ability to fully capture the composition of the host microbiome. However, high-throughput sequencing is a promising approach better able to characterize human microbiome function and composition. Such characterization is essential in determining the role microbiota play in disease development, especially when comparing microbiomes of healthy and disease subjects. Recently, an increasing number of sequencing studies offer enormous possibilities to study microbiome functionality and composition across different individuals and diseases. Current approaches use existing reference databases to classify and identify microbial communities present in the individual host, and then compare these classifications. However, existing databases are far from complete, and rely only on a limited compendium of reference genomes, thereby limiting the ability to accurately determine microbial compositions. This further confounds cross-subject microbiome comparisons. Ideally, one would be able to use all the microbial reads to determine patterns of diversity in health and disease, not just those that classify to existing databases.

In this paper, we use a reference-free comparison method on meta-transcriptomics data to characterize microbial compositions across healthy and disease samples. We use publicly available RNA-Seq datasets from a coronary artery calcification study [23] of eight cases and eight controls (matched for gender, age and ancestry) to detect microbial communities and perform species independent comparisons. Reads that failed to map to the human genome are used as nonhost sequencing reads to detect the presence of microbes. To capture both differences in microbiota abundances and composition we propose a reference-free approach (EMDeBruijn) able to characterize individual metagenomic samples based on an associated de Bruijn graph (that is, a certain directed graph on *k*-mers). This approach provides a powerful, species independent way to assess microbial diversity across individuals and subjects. Incompleteness of existing microbial databases becomes a non issue as sequencing information is translated directly into a de Bruijn graph. Properties of the de Bruijn graphs are then used to compare microbiome composition across individuals. Briefly, this new metric measures the minimal cost of transforming one sample's associated de Bruijn graph into another one to measure the similarity of the microbial communities between two samples. This reduces to using the Earth Mover’s Distance (EMD), and we utilize a recent implementation of the EMD that takes advantage of the underlying graph structure. We call our method *EMDeBruijn*. Code implementing this method is available at github.com/dkoslicki/EMDeBruijn.

The ability to determine the association of the microbial communities with different types of diseases may have implications for disease diagnoses and prevention. Detection of any disease-promoting effects of microbiota for the different types of diseases may suggest the necessity to routinely incorporate the meta-omics analysis in the current genetics-based studies of the diseases [21]. Additionally, accurate characterization of the personal microbiome and ability to compare it across individuals and diseases opens new possibilities to investigate host-microbiome interactions.

## 2. METHOD

### 2.1 Overview

Words of length *k* on a given alphabet 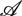, also known as *k*-mers, are used extensively in genomics problems. While reads from next-generation sequencing technologies may be short, the dimensionality of all, say, 6–mers on the alphabet 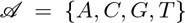is much smaller than the dimensionality of all strings of length ~ 100nt on 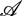. Furthermore, the counts or frequencies of all *k*-mers in a given dataset can give an analytically tractable “signature” associated to a dataset. Many *k*-mer based metagenomic classification methods use a variety of metrics when comparing such count or frequency vectors [12, 20, 26]. Many metrics are used to compare these vectors, and there has been no consensus as to which metric/metrics are best suited for the task. A further problem arises due to the fact that when attempting to analyze metagenomic data, samples are typically classified via some approach (either *k*-mer based [4,12,13,20,26] or non-*k*-mer based [15]), and then further analysis (clustering, PCoA, etc.) is performed using a variety of metrics on the vectors of relative species abundances. It has been shown [11] that the selection of the metric (typical examples include Euclidean distance, Jensen-Shannon divergence, Bray-Curtis, weighted/unweighted UniFrac metrics) significantly affects downstream analysis (such as determining en-terotypes across the human body).

Comparing samples by using a metric on the vector of classified relative abundances has a number of drawbacks. Beyond misclassifications and unclassified portions of the sample (which can comprise a very large percentage of a given sample), this approach is constrained by the completeness and correctness of the training database utilized. It is well-known that current bacterial databases suffer from mislabeled bacterial sequences and in general only represent a very small percentage of predicted bacterial species.

We present an approach that aims to resolve two of these issues: first, we develop a reference-free distance metric between metagenomic samples that is able to accurately characterize the similarities/differences between samples that bypasses the need to first classify the samples. Second, this metric takes advantage of the de Bruijn graph structure of *k*-mer frequency vectors and can be shown mathematically to be a natural choice when comparing *k*-mer frequency vectors.

The usage of de Bruijn graphs in bioinformatics is practically ubiquitous. Some methods have been developed to utilize de Bruijn graphs for taxonomic profiling, but still rely on a known database of organisms [3, pg. 11] and so are not entirely reference free. Other problems, such as genome assembly, rely almost exclusively on properties of de Bruijn graphs [7, 14, 27] or overlap graphs. However, there exist many aspects of de Bruijn graphs that have yet to be exploited.

One such aspect is the fact that de Bruijn graphs allow for the development of a rather natural metric on *k*-mers. The metric we propose to use is the Earth Mover’s Distance (also known as the first Mallows distance, first Wasserstein distance, etc) between *k*-mer frequency vectors on the de Bruijn graph of order k on the alphabet {A, C, G, T}.

**Figure 1:**
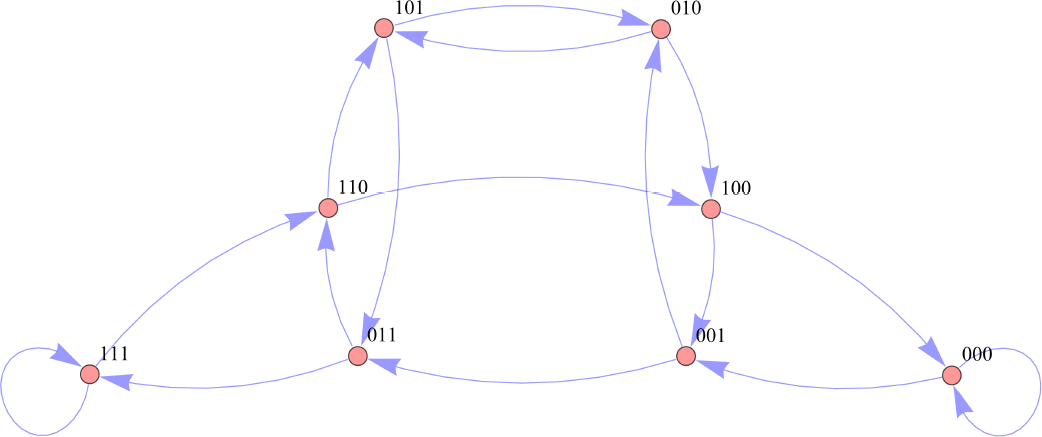
Graph of *B*_3_ ({0,1}).

### 2.2 Earth Mover's Distance on de Bruijn graphs (EMDeBruijn)

We describe here in detail the reference-free method we utilize to measure the differences between individual meta-transcriptomic samples, called *EMDeBruijn,* and first set some notation. Let 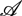 be an alphabet of finitely many symbols. Typically 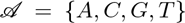. For 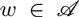, let |*w*| denote the length of *w* (or equivalently, the total occurrence of each symbol of 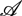 in *w*). For k ϵ 葩, let 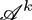 denote the set of all words *w* with |*w*| = *k*. Let 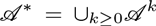 be the set of all finite length words on 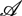. Write the letters of *w* = *w*_1_*w*_2_…*w*_k_ for |*w*| = *k*. For *w*, 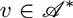, let *occ_v_*(*w*) = and freq_*v*_ (*w*) be the number of occurrences and frequency of occurrence respectively of v in w with overlaps. I.e.

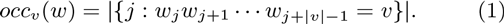

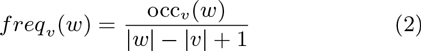

By freq*_k_*(*w*) we mean the vector 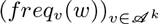. We are now in a position to define the de Bruijn graph and the EMDeBruijn metric.

#### Definition 1.

(*de Bruijn graph*) Denote B_k_(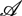) to be the graph with vertex set V = A^k^ and edge set

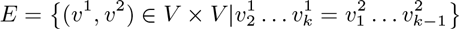

Note that this means an arrow is drawn between two vertices if the suffix of the first vertex matches the prefix of the second vertex. We now consider the undirected graph.

#### Definition 2.

(*Undirected de Bruijn graph*) Let 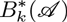 to be the graph with vertex set V = 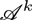 and edge set

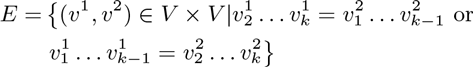

The graph of B3({0, 1}) is given in figure 1, and a graph of 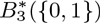 is given in figure 2. Next, we need to define a *flow* (also called *coupling*) between *k*-mer frequency vectors. A visualization of a flow is given in figure 3.

**Figure 2:**
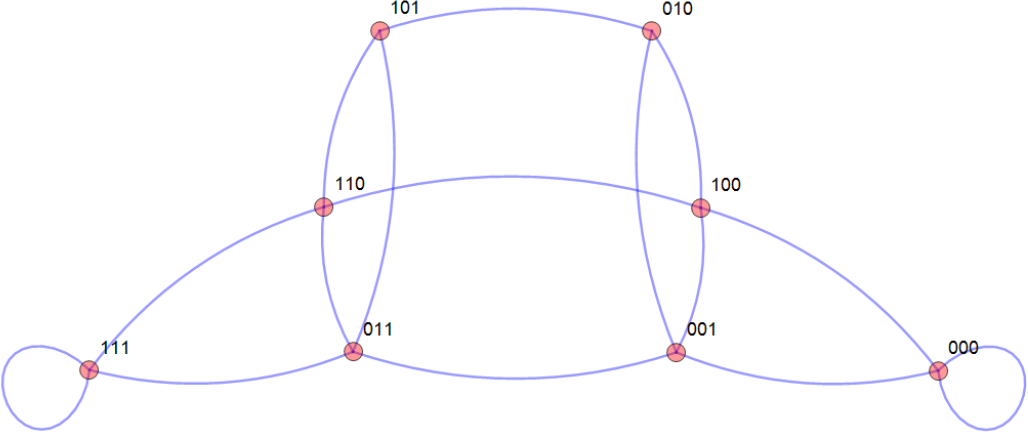
Graph of 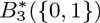

#### Definition 3.

*(*Flow/Coupling*) For *w*, 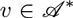and *k* ϵ 葩, a flow (or coupling) of *freq*_k_*(*w*) *freq*_k_(*v*) is a matrix 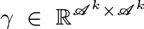 such that 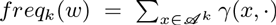 and 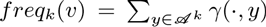. Let Γ(*w*,*v*) denote the set of all couplings of *w* and *v*.

We are now in a position to define the metric that will comprise out method. For the following, recall that for connected graphs such as these, the graph distance between two vertices is the shortest path in that graph between the two vertices.

#### Definition 4.

*EMDeBruijn* Let 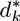be the graph distance associated to 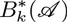. Then

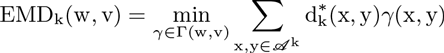

Since the set of couplings Γ(*w*,*v*) depends only on the frequencies of *k*-mers in *w* and *v*, definition 4 can take as inputs the *k*-mer frequency vectors of two samples consisting of multiple sequences. Hence, the metric in definition 4 measures the minimal cost (in terms of distance) of transforming one sample’s *k*-mer frequency vector into the other sample’s *k*-mer frequency vector where transformations are restricted to moves along the de Bruijn graph.

To compute the EMDeBruijn metric in definition 4, we use the FastEMD [18,19] implementation of the Earth Mover’s Distance since the graph metric is naturally thresholded to the diameter of the de Bruijn graph. This reduces to using a min-cost-flow algorithm to solve the optimization problem in definition 4. When counting *k*-mers, we always include the original sequence, as well as its complement. Again, the advantage of definition 4 is twofold: first, it bypasses the need to first classify a sample, and secondly, it takes advantage of the natural de Bruijn graph structure of *k*-mers (instead of treating *k*-mers as independent of each other). The current implementation of EMDeBruijn has a runtime which is quadratic in the number of *k*-mers (and so exponential in the *k*-mer size). Further algorithmic improvements to reduce this computational complexity are currently being developed, but at the moment we restrict the number of *k*-mers to be less than or equal to 4,096 and hence consider k = 6 (resulting in a total running time of approximately 55 CPU minutes). A heuristic algorithm that allows for faster execution time and increased *k*-mer size (at the expense of non-optimal results) is currently being developed.

## 3. RESULTS

### 3.1. Blood microbiome profiling across CAC and non-CAC patients

We study the microbiome composition and abundance levels of the microbial communities present in the blood across coronary artery calcification (CAC) patients and controls. The framework of the study is summarized in figure 4. Publicly available RNA-Seq data was obtained from a CAC study [23]. The data consisted of the peripheral blood Illumina RNA-Seq samples from eight cases and eight controls matched for gender, age and ancestry. Peripheral blood was collected using the PAXgene RNA tubes and the PAXgene Blood RNA Kit IVD. Sequencing was performed on Illumina GAIIx sequencers generating paired end sequencing 2X76bp. First we extract non-host reads from each sample by mapping reads onto the human genome. An average of 68% of the reads of each sample aligned to the human hg19 reference genome via tophat2 (v2.0.4) [10] with the default settings. Additionally, tophat2 was supplied with the gene annotations to improve the quality of the mapping. Non-host reads (reads failing to map to the human reference genome) are used as a candidate microbial reads to detect the presence of microbial communities and compare them across host individuals and subjects. Microbial reads are expected to be a small portion of the non-host reads due to intensive immune system response able to deactivate the majority of bacteria, fungi, viruses and parasites in the blood. Besides the microbial reads, non-host reads contain low-quality and/or low complexity reads as well as reads from repetitive elements, circular RNAs (circRNAs), gene fusion events, trans-splicing events, recombined B and T cell receptor (BCR and TCR) loci [16]. Reads containing larger number of sequencing errors are more likely to be ignored by the current mapping algorithms. We perform additional Megablast [1] alignment of the non-host reads against the human reference to exclude any reads ignored by the alignment tools.

The filtered set of non-host reads is used to determine the composition of the blood microbiome. We use a novel reference-independent approach able to perform a species independent comparison of the microbial communities, and refer to this method as EMDeBruijn. This method uses directly the non-host reads and condenses them into a de Bruijn graph (whose vertices are *k*-mers). We then use the Earth Mover’s Distance (EMD) metric on this graph to measure the differences between the samples based on the putative microbial communities.

Additionally, we examine any potential experimental contamination (contamination by the microorganisms introduced during the sequencing experiment). We use additional RNA-Seq samples from the The Cancer Genome Atlas (TCGA) [25] to confirm our finding of a microbial presence in blood samples, and examine the possibility of experimental contamination.

### 3.2 Blood microbiome contains known reference microbial genomes and rRNA genes

We confirmed a microbial presence in both the control and case samples by using standard community profiling techniques. For whole meta-transcriptome classification, we used MetaPhlAn [22] version 1.7.7 with bowtie2 [8] version 2.2.3 to classify the non-host reads. Using the default settings, we are able to successfully classify an average of 0.06% of the non-host reads (~ 24K reads per sample).

We further confirm this result by extracting the 16S ribo-somal sequences present in the non-host reads. We selected the non-host reads having an exact match to at least one of 22 “universal” 16S rRNA primers [24]. On average, 145K reads were extracted per sample. Extracted reads were then classified by the Ribosomal Database Project’s (RDP) classifier [26] using version 2.8 and the default settings. On average, only 1% of the reads were classified to the genus level.

**Figure 3:**
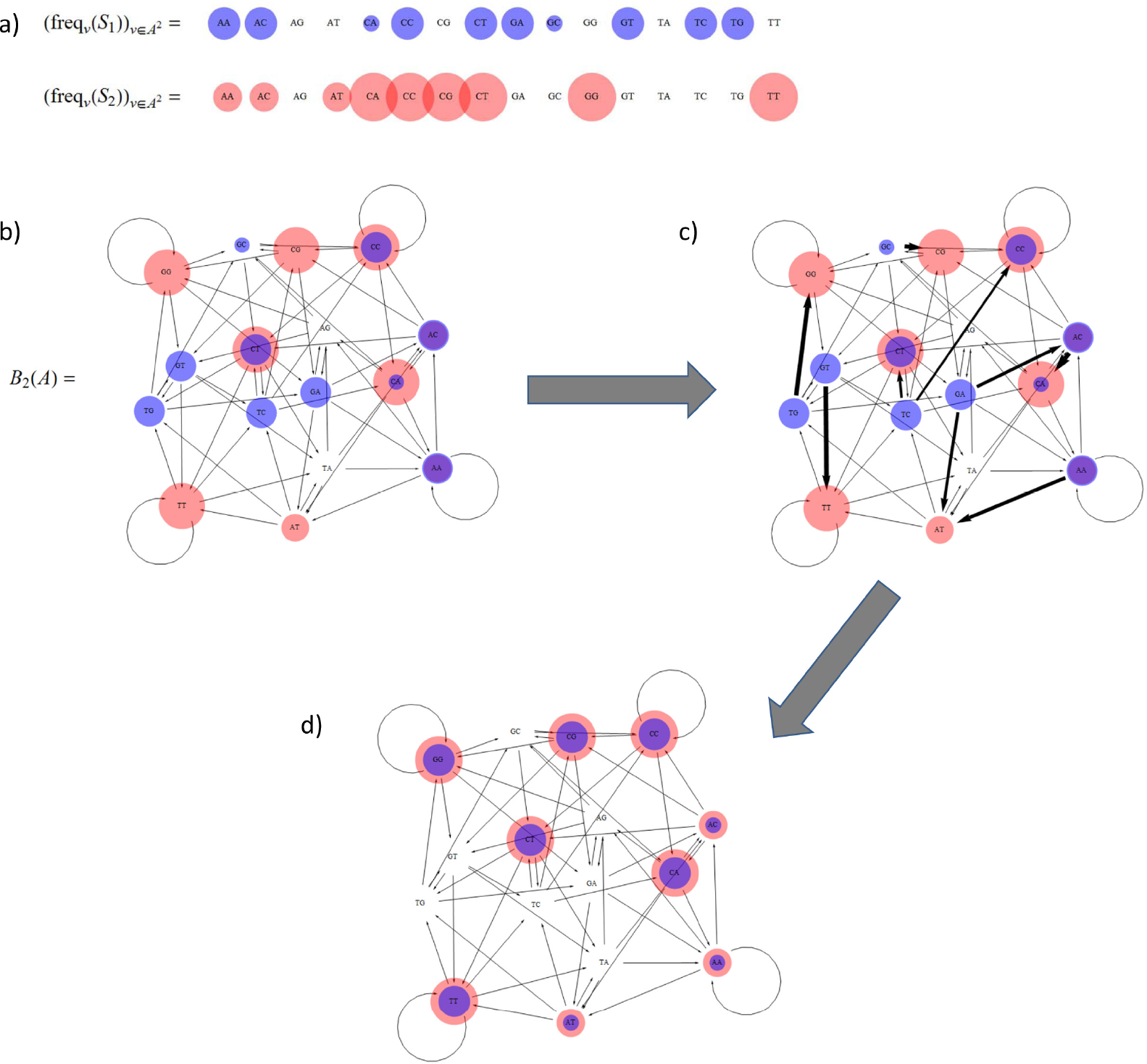
Visualization of the EMDeBruijn Distance. a) Pictorial representation of 2–mer frequencies for two hypothetical samples, S_1_ and S_2_. b) The 2–mer frequencies overlaid the de Bruijn graph 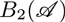. c) Representation of the ow used to compute EMD2(S_1_; S_2_); dark arrows denote mass moved from the initial node to the terminal node. d) Result of applying the flow to the 2–mer frequencies of S_1_.

Next we evaluate microbial community variability across the CAC patients and controls. The classified blood microbial communities were observed to vary across the samples, however no disease-specific patterns were observed (Figures 5 and 6). The clear discrepancy between the MetaPhlAn and the 16S rRNA approach (RDP's NBC) indicates the disadvantage of using these approaches, suggesting that a reference-free approach is required to account for all the organisms, including ones not present in any of the databases.

To confirm the blood taxonomic composition in the CAC study and to examine any hypothetical contamination we use seven whole blood RNA-Seq samples from the TCGA collected from the Acute Myeloid Leukemia (LAML) patients (see Appendix section A.2). A total of approximately 18% and 47% of the genera in the CAC samples were also observed in the LAML samples using MetaPhlAn and RDP’s NBC respectively. Furthermore, these shared genera contributed an average of approximately 37% and 51% of the relative abundance in the CAC samples as measured by MetaPhlAn and RDP’s NBC respectively. The presence of similar genera of bacteria across different studies and experimental protocols corroborates the claim that a bacterial presence is genuinely being observed in the given samples (and is not due to sequencing artifacts or contamination).

To compare the CAC patients to the controls, we created a PCoA plot using the Jensen-Shannon divergence on the genera level reconstructions for both the MetaPhlAn and RDP reconstructions. The results are given in figure 6. Neither MetaPhlAn nor RDP’s NBC are able to effectively distinguish between case and control samples. Other distance metrics were also employed (results not shown) with the same qualitative outcome.

To quantify how well MetaPhlAn and RDP’s NBC distinguished between cases and controls, we performed a hierarchical clustering using the Jensen-Shannon divergence on the genus level reconstructions of the two methods. We took the top two resulting clusters (which consisted predominantly of either case samples or control samples). Treating these as classifiers, we obtained the classification measures contained in (table 1. Dendrograms and heatmaps of these results are also contained in the appendix. We performed a similar analysis using UPGMA trees (results in the appendix section A.4).

**Figure 4:**
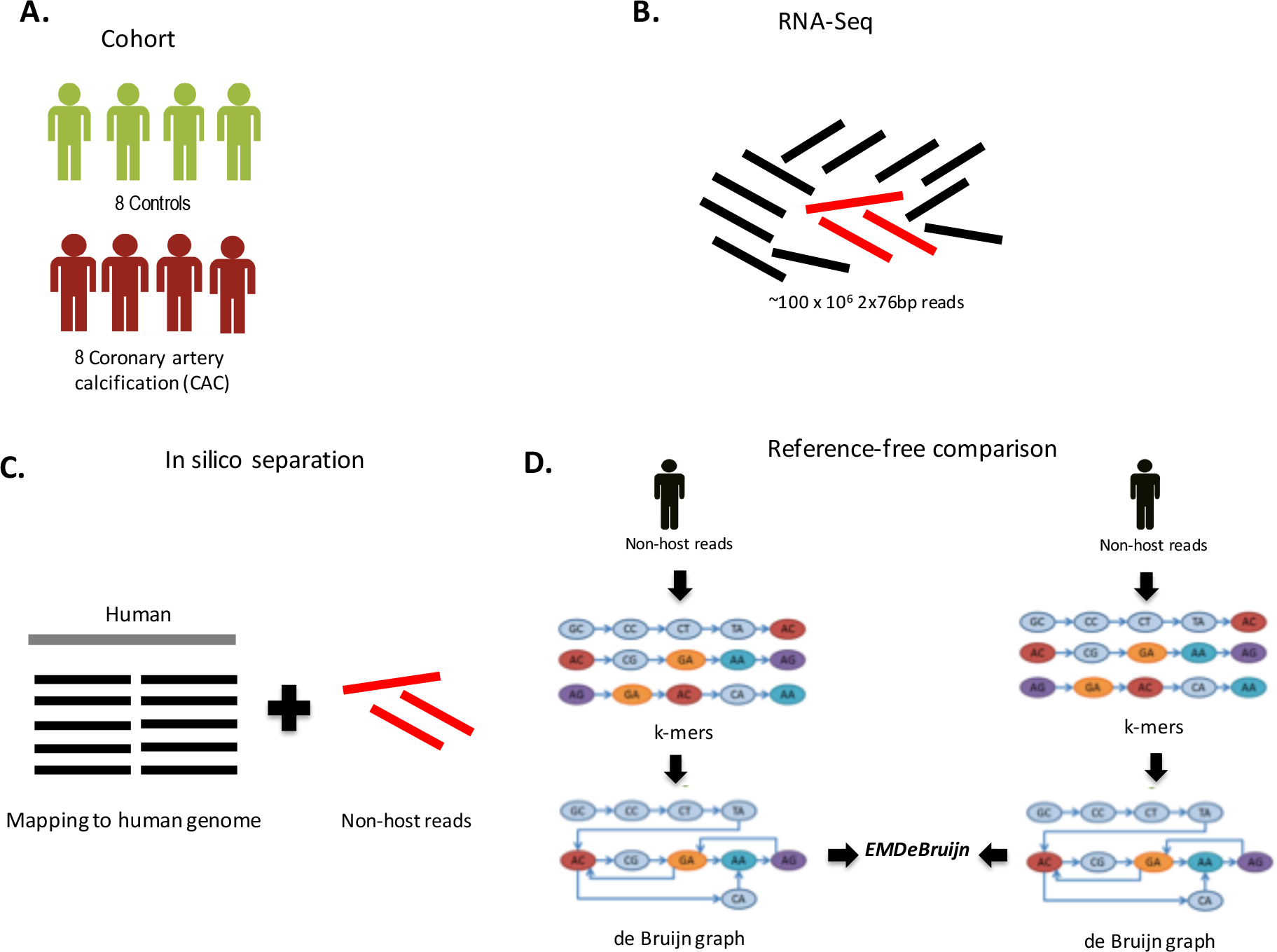
Framework of the study. (a) We analyze cohort of eight cases and eight controls matched for gender, age and ancestry, (b) Sequencing was performed on Illumina GAIIx sequencers generating paired end sequencing 2X76bp. (c) Nonhost reads (reads failing to map to the human reference genome) from each sample were extracted by mapping reads onto the human genome. Non-host reads are used as a candidate microbial reads to detect the presence of microbial communities and compare them across host individuals and diseases, (d) Non-host reads are used to determine the composition of the blood microbiome. We use a novel reference-independent approach (EMDeBruijn) able to perform a species independent comparison of the microbial communities. EMDeBruijn condenses the reads into the de Bruijn graph. The vertices of the de Bruijn graph are fc-mers produced from the sequencing reads. Earth Mover’s Distance (EMD) metric is used to measure the differences between the samples based on the proprieties of the graph.

**Figure 5:**
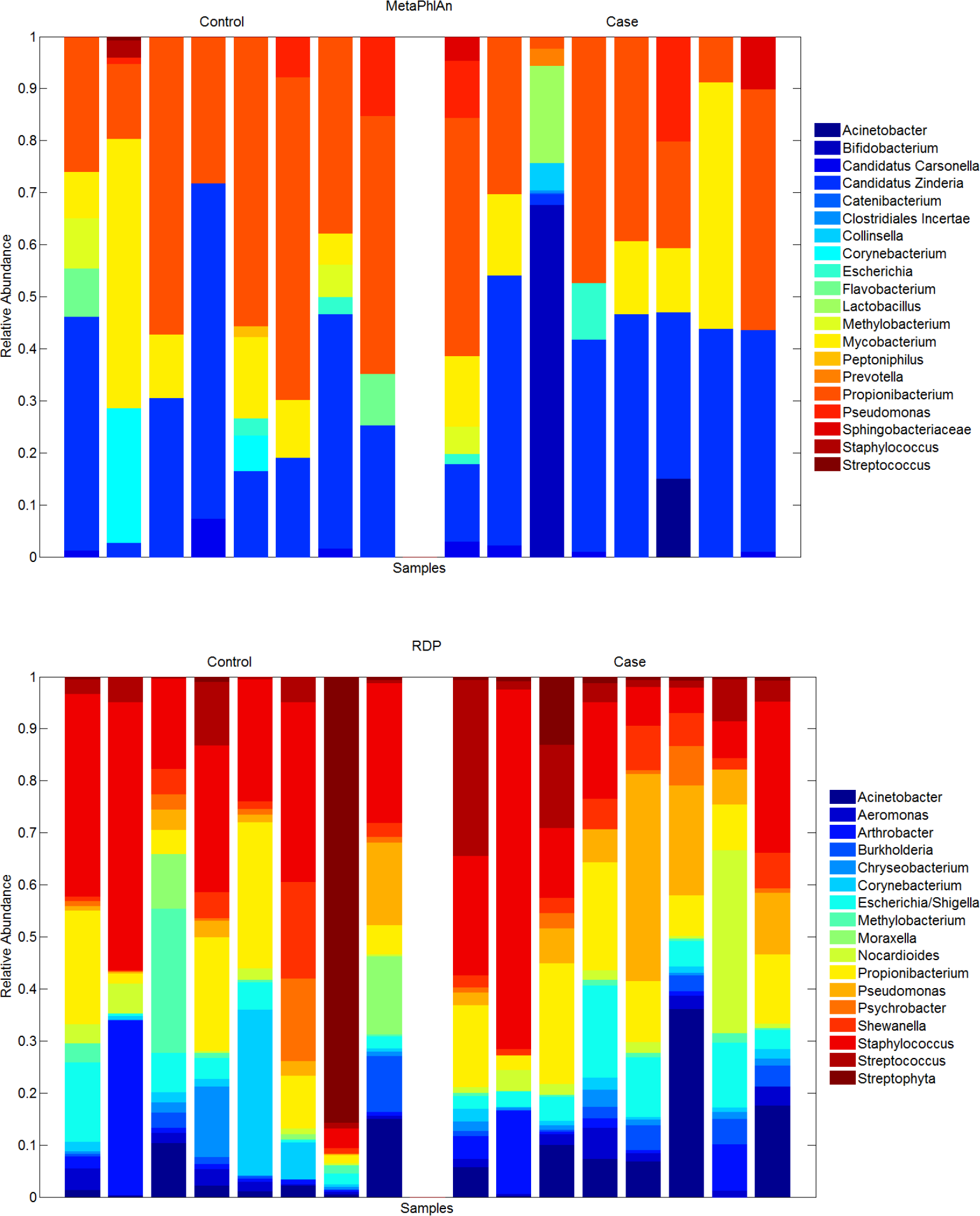
Relative abundances of blood microbial communities across all subjects. Non-host reads were classified at the genus level.

Clearly, available microbial reference databases allow one to classify only a small portion of non-host reads, thus biasing any further analysis of the microbiome composition. We propose a reference-free approach that is able to use all the putative microbial reads to characterize the microbial communities and compare them across subjects.

**Figure 6:**
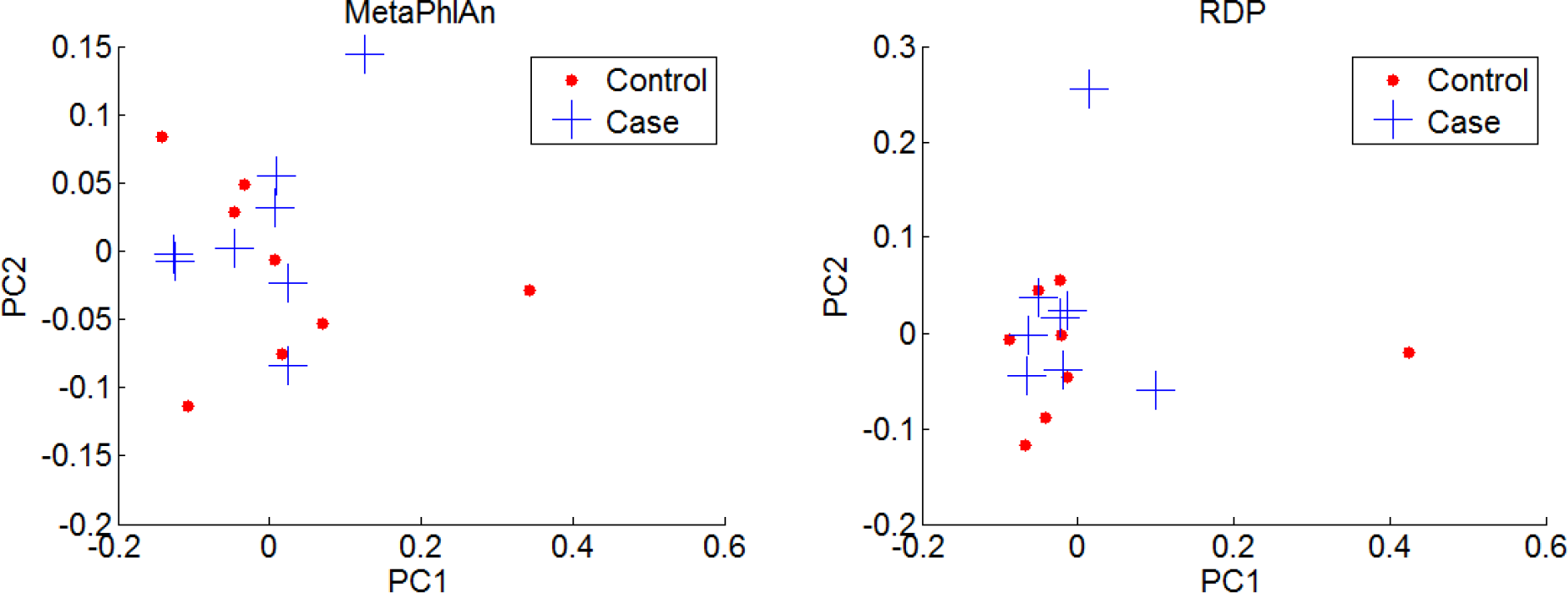
PCoA plots using Jensen-Shannon divergence on the genus level reconstructions.

**Figure 7:**
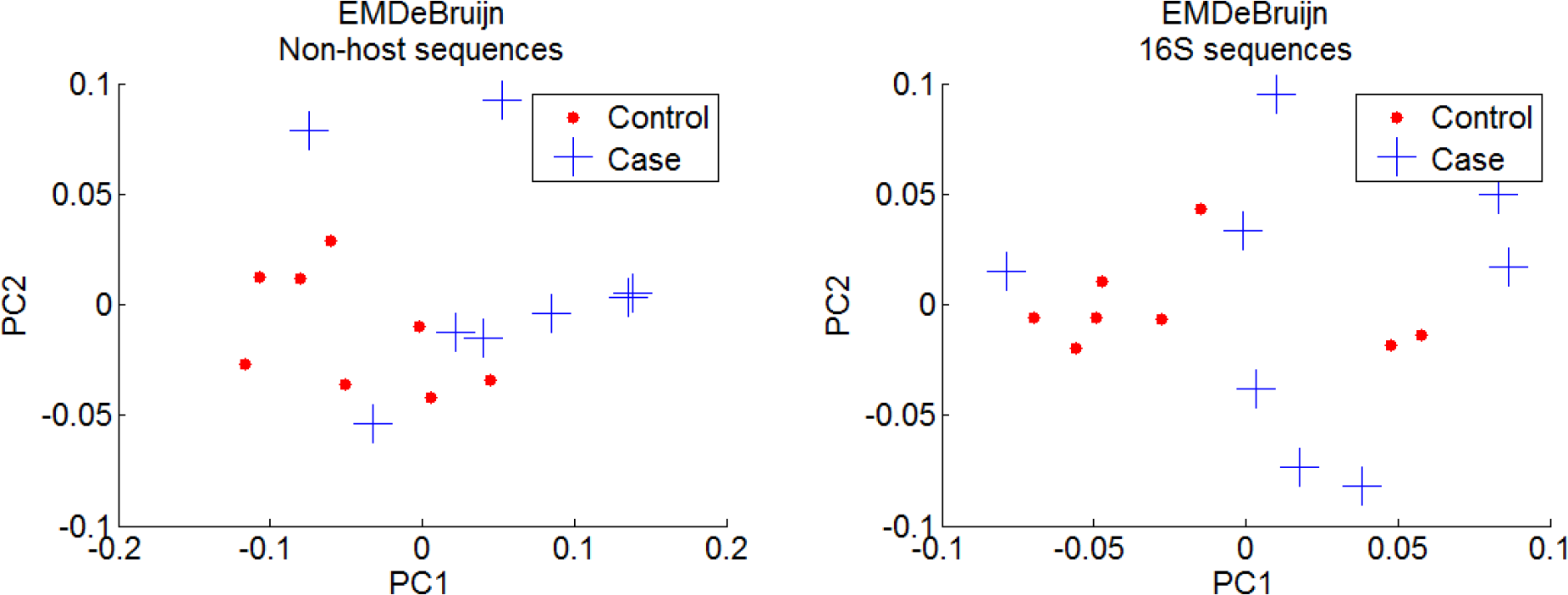
PCoA plot using the EMDeBruijn metric. Note the clustering of the control samples.

### 3.3 Reference-free comparison of microbial communities across CAC and non-CAC patients

Reference-based microbiome profiling is limited to discover the microbial taxa present in the available reference databases, which are known to be far from complete. This limits the ability to accurately profile microbial communities and perform cross-individual microbiome comparisons. Profiling performed by reference-based methods (Section 3.2) are inconsistent, and have limited ability to discriminate between healthy from disease samples. We argue that comparing the taxonomic classifications performed both by MetaPhlAn and RDP have a limited possibility to discriminate the samples into a health and disease group. We propose an alternative approach able to overcome the incompleteness of existing microbial databases by directly using the sequencing information condensed into a de Bruijn graph. Applying the proposed EMDeBruijn method (Section 2) for the *k*-mer size of k = 6, we obtain a metric which is applied to discriminate between healthy and disease samples. The EMDeBruijn metric as defined in definition 4 is used to produce a PCoA plot (Figure 7). This method is able to clearly group the control CAC samples, suggesting a possible disease promoting effect of the microbial communities yet to be classified.

As with the reference-based techniques, we performed a hierarchical clustering using the EMDeBruijn metric. After performing the clustering, we took the top two clusters (which consisted predominantly of either case samples or control samples). This effectively partitioned the data, which we then treated as a classifier. We then obtained the classification measures contained in (table 1. Dendrograms and heatmaps of these results are also contained in the appendix (see figure 13). We performed a similar analysis using UPGMA clustering, the results of which were identical to the hierarchical clustering. These results are contained in the appendix section A.4. In all, this demonstrates that the case and control sample are most effectively distinguished when using the EMDeBruijn metric.

Whether the reads driving the differences between the health and disease are microbial is an open question. To address this question, we perform the species independent comparison of the microbial communities across the samples based on the 16S ribosomal sequences extracted from the non-host reads. Ribosomal sequences are specific to microbial organism allowing to confidently extract the reads corresponding to the microbial communities. Results on the 16S reads (see figure 7) suggest similar discrimination of the samples.

**Table 1:**
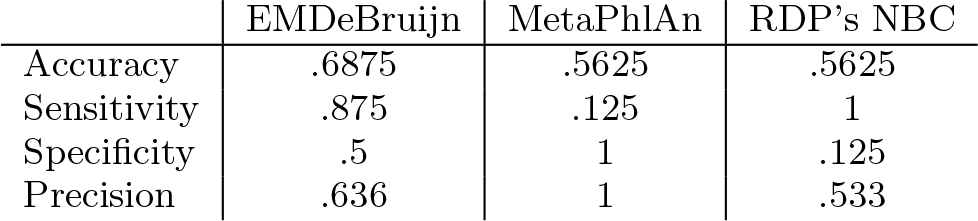
Classification results for the hierarchical clustering derived from the three classification methods. These measures result from selecting the top/largest two clusters.

As evidence that the EMDeBruijn metric is a more useful *k*-mer based metric than other commonly utilized metrics (such as the Jensen-Shannon or Kullback-Leibler divergence), we created PCoA plots using a number of these metrics directly on the non-host 6–mer counts and 10–mer counts. As shown in Appendix section A.1, figure 8, and section A.4, none of these methods clusters as well as the EMDeBruijn metric. Additionally, we check the classification power of using *k*-mer frequencies alone on the CAC vs non-CAC samples. We observe similar frequencies of 6–mers in both healthy and disease groups. The two 6–mers with frequencies higher in the control group than the case group were CCCCCC and GGGGGG (Appendix 14).

To investigate the effect of the human reads on the ability to differentiate between the healthy and disease samples, we applied our method for all the sequencing reads. We observe similar discrimination into health and disease suggesting that adding human sequences provides no improvement (see Appendix section A.3).

**Figure 8:**
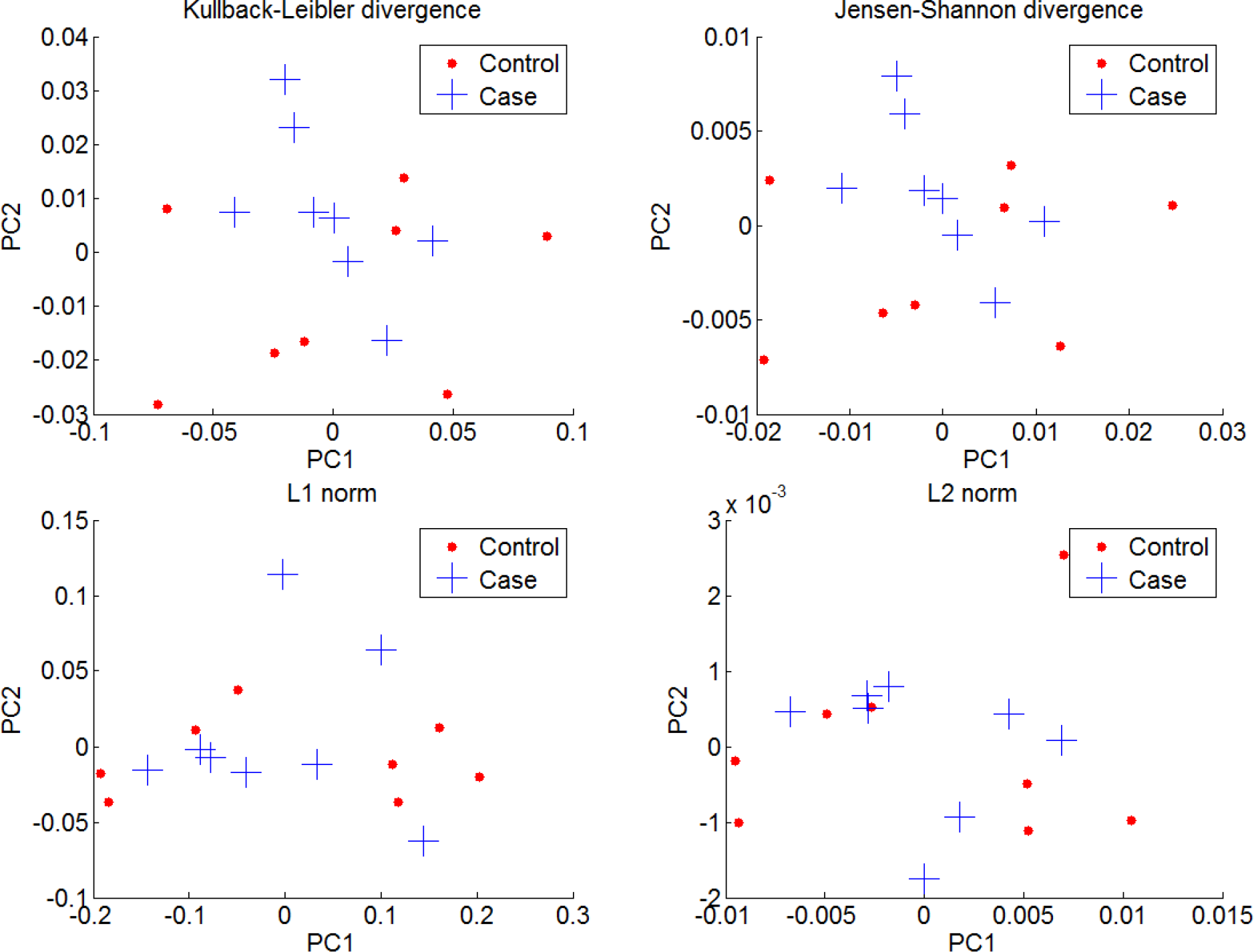
PCoA plots corresponding to four different metrics applied directly to the 6–mer counts.

One of the current limitations of this method is the absence of a straightforward approach to extract the taxa driving the difference between the samples. One way to overcome this limitations is to extract the sequences contributing to the differences between the samples and match those against the hypervariable taxa-specific gene families. Hypervariable regions from gene families are previously identified to be as nearly universal, allowing one to differentiate between species and taxa [5]. However, a large number of sequences are observed to contribute to the differences between the CAC and non-CAC samples. Further investigation is required to quantify the contribution of different taxa to the observed case and control clustering.

## 4. DISCUSSION

In summary, meta-transcriptomics profiling was used to determine the composition of the blood microbiome across coronary artery calcification (CAC) patients and controls in an effort to determine the relationship between the blood microbiome and CAC disease.

First, we seek to determine the microbial composition of the blood across CAC and control patients. We use non-host RNA-Seq reads to perform the taxonomic classification using existing computational methods. Both MetaPhlAn and RDP’s NBC were able to discover various microbial communities across the health and control samples. However neither of these methods were able to find any disease-specific patterns in the microbiome nor were able to discriminate the samples into disease and healthy groups. Furthermore, the genera level classification provided by both methods shows large discrepancies. This reveals the limitations of these methods, namely relying on the known microbiome databases to classify the metatranscriptomics samples. One way to overcome this limitation is to directly use the sequencing data to determine patterns of diversity in health and disease, thereby avoiding the bias introduced by the existing databases.

We then proposed a novel EMDeBruijn approach which provides a powerful reference-independent way to assess microbial diversity across the samples. It allows one to condense the sequencing data into a de Bruijn graph. We use Earth Mover's Distance (EMD) to measure the similarities of the microbial communities via their associated de Bruijn graphs. The ability to account for all the candidate microbial reads allows our method to captures information relevant to the disease and differentiates between the case and control CAC samples. All this suggests that a reference-free approaches is a preferable choice in comparative metatranscriptomics studies.

In de Bruijn-based approaches one of the most significant parameters is the fc-mer size. Longer fc-mer size is usually preferable for studies performing genome assembly, as this provides the possibility to bridge repetitive genomic regions. In this study we do not aim to assemble the microbial genomes and instead measure the similarities and differences of the microbial communities across different conditions. While the full set of *k*-mers is present when using *k* = 6, the variability of bacterial sequences results in *k*-mer frequency profiles whose difference can be detected by EMDeBruijn. It is reasonable to assume that any contribution of unmapped human-related sequences to the *k*-mer frequency profiles is relatively constant across the samples (especially when using shorter *k*-mer sizes).

Altogether, this study of microbial communities suggests an important role of the microbiome in CAC disease and indicates the presence of the disease-specific microbial community structure in the CAC patients. Establishing the causal relationship between the microbiome and CAC disease is yet to be studied and requires additional inquiry.

## A. SUPPLEMENTARY MATERIAL

This appendix contain additional material supporting the main text.

## A.1 Other *k*-mer based metrics

We demonstrate here that the EMDeBruijn metric is more effective at extracting relevant information than more traditional metrics applied directly on the *k*-mer counts. To that end, we formed the 6mer counts for each of the non-host CAC data sets. We then created PCoA plots corresponding to a variety of commonly utilized metrics. The results are shown in figure 8. Note that none of these metrics give nearly as clear a clustering of the control samples as the EMDeBruijn metric does.

To verify that this is lack of clear clustering is not due to the relatively small *k*-mer size of *k* = 6, we repeated the same process as before, but this time for *k* = 10. As seen in figure 9, the increased *k*-mer size only marginally qualitatively improves the clustering. This can be quantitatively confirmed, as seen in Table 3, where it is shown that the classification metrics when *k* = 10 are still worse than when using *k* = 6 and EMDeBruijn.

**Figure 9:**
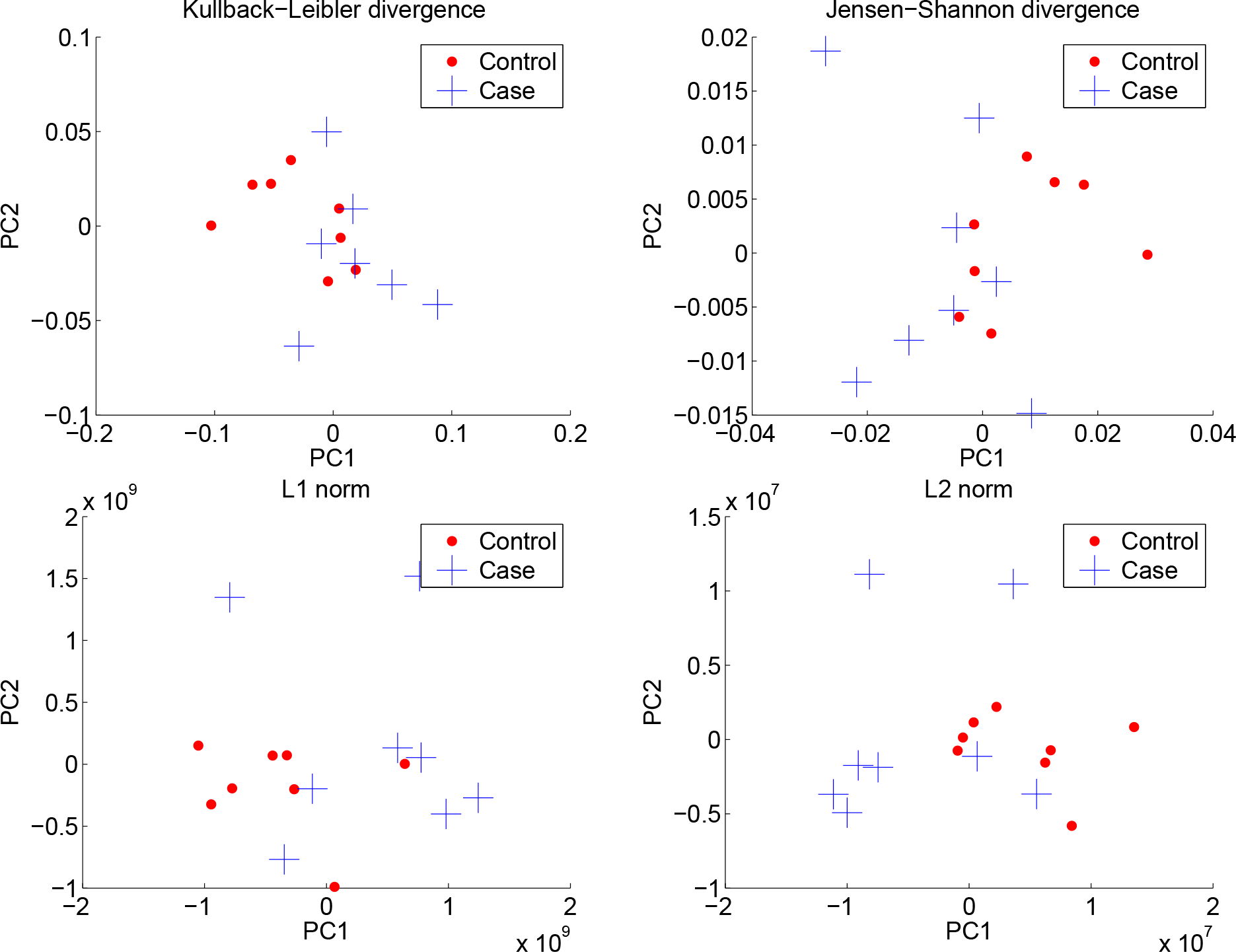
PCoA plots corresponding to four different metrics applied directly to the 10–mer counts.

## A.2 Comparison to LAML and colorectal data

To confirm the blood taxonomic composition in the CAC study and to examine any hypothetical contamination we use seven whole blood RNA-Seq samples from the TCGA collected from the Acute Myeloid Leukemia (LAML) patients. The LAML data was obtained from 173 RNA-Seq primary blood derived cancer samples available from the The Cancer Genome Atlas (TCGA) from CGHUB at the USC. We randomly selected 7 samples to determine their microbial content. The same procedure detailed in section 3.2 was utilized to extract non-mapped reads and classify them via MetaPhlAn and further extract 16S reads and classify them via RDP’s NBC. An average of 75% of the reads aligned to the human genome, and of these, 0.5% were classified by MetaPhlAn and 7.7% of the extracted 16S rRNA sequences were classified down to the genus level using RDP’s NBC. As we are concerned with the shared common genera, we restricted our attention to only those genera whose abundance in either the CAC or LAML data was greater than 5% when summed over all samples. Included in figure 10 are Venn diagrams representing the overlap between the classified genera between these LAML samples and the CAC samples. A total of approximately 18% and 47% of the genera in the CAC samples were also observed in the LAML samples using MetaPhlAn and RDP’s NBC respectively. The presence of similar genera of bacteria across these different studies supports the claim that a bacterial presence is genuinely being observed in the given samples.

**Figure 10:**
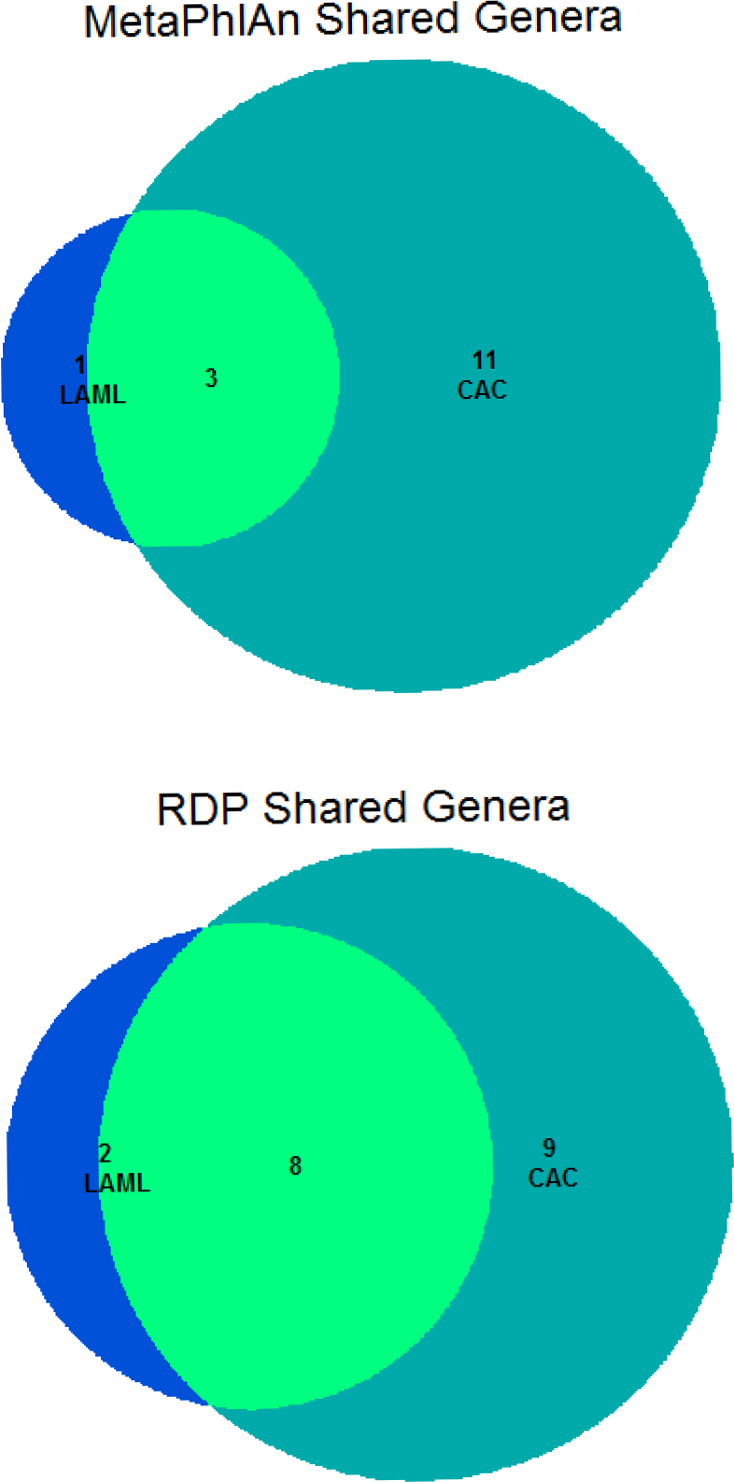
Number of genera shared between LAML and CAC data sets.

Another important observation is that using RNA-Seq allows for the sequencing of non-host genomic material. To demonstrate this, we also performed the same analysis procedure detailed in section 3.2 on 9 whole genome shotgun primary colorectal adenocarcinoma samples, along with 9 whole genome shotgun controls samples [2]. Analyzing this data revealed no presence of any known microbial communities. This suggests that RNA-Seq may be the technology of choice for profiling microbial communities in human tissues when there is an expected minor microbial presence.

## A.3 Utilizing host and non-host reads

To investigate the effect of the human reads on the ability to differentiate between the healthy and disease samples, we utilized the EMDeBruijn method on all the raw sequencing data. Figure 11 shows the resulting PCoA plot. Note the similar clustering of the control samples as in figure 7. This indicates that the addition of the mapped reads does not seem to significantly influence the case/control clustering as seen in figure 7.

**Figure 11:**
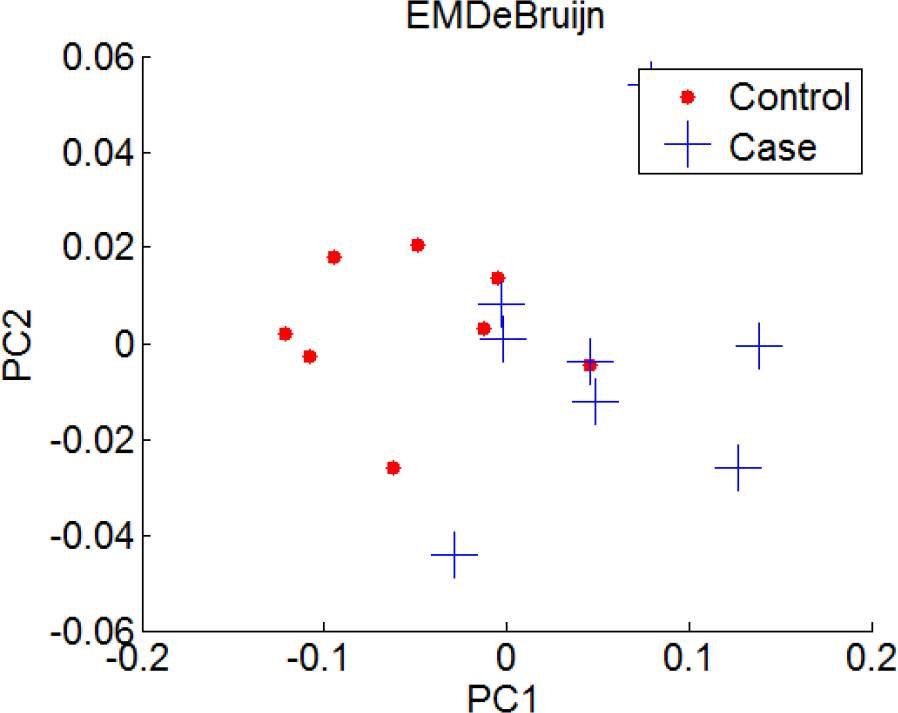
PCoA plot using the EMDeBruiujn metric on all the CAC sequences (both host and non-host).

## A.4 Verification of clustering accuracy

Having already supported the claim that MetaPhlAn and RDP’s NBC do not distinguish between case and control samples as effectively as EMDeBruijn by using hierarchical clustering (see table 1 and section 3.2), we further confirm this by using UPGMA clustering as well. Hence, we built UPGMA trees by using the Jensen-Shannon divergence on the genus level reconstructions of RDP's NBC and MetaPhlAn. We also used the EMDeBruijn metric to form a UPGMA tree as well. We took the top two clusters (which consisted predominantly of either case samples or control samples) and treated them as a classifier. We obtained the classification metrics contained in (table 2. Visualizations of the UPGMA clustering is contained in figure 12.

**Figure 12:**
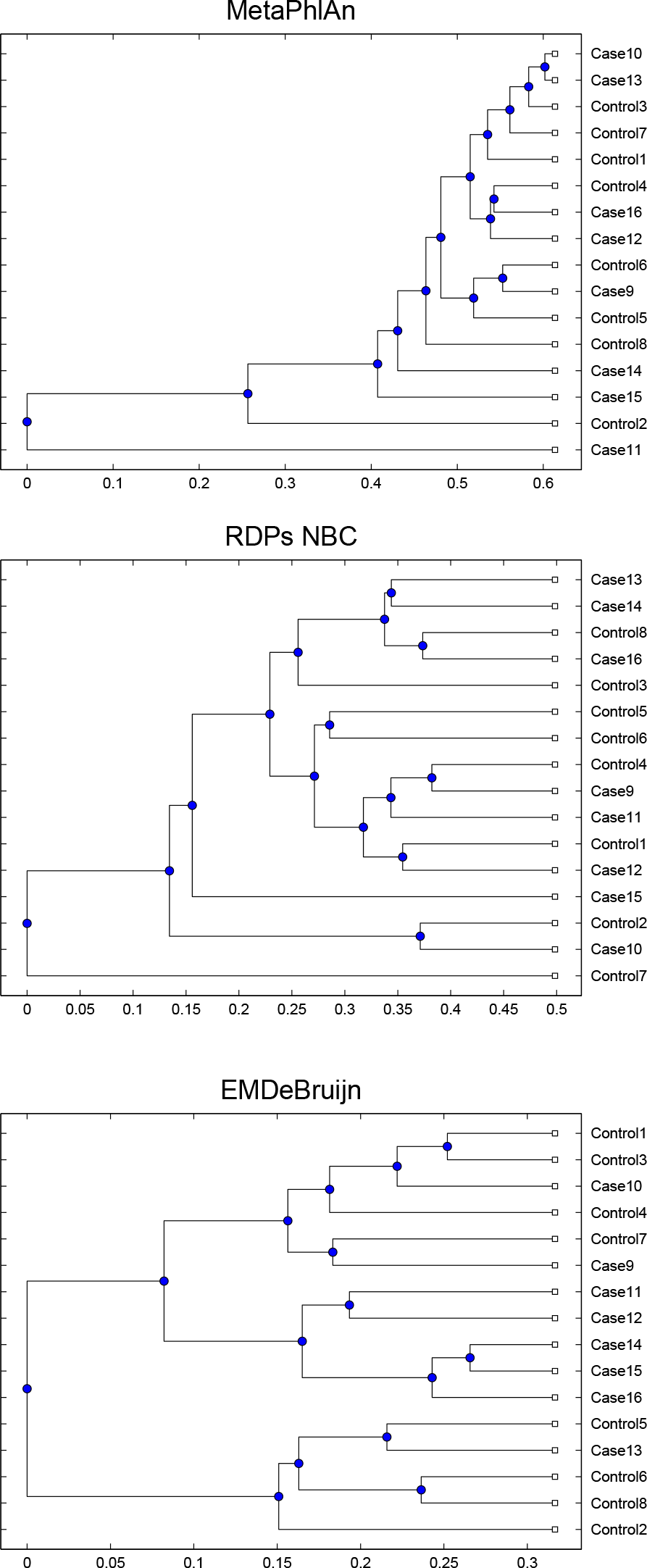
UPGMA trees based on the Jensen-Shannon divergence of the genus level classification for MetaPhlAn and RDP’s NBC, as well as the UPGMA tree based on the EMDeBruijn distance metric on the non-host reads.

Furthermore, we formed UPGMA trees for the reference-free metrics as well. To demonstrate that a smaller *k*-mer size for EMDeBruijn is still more effective than using a larger *k*-mer size for the other methods, we used the *k*-mer size of *k* = 6 for EMDeBruijn, and *k* = 10 for the Jensen-Shannon divergence, Kullback-Leibler divergence, L1 norm, and L2 norm. The results are contained in table 3 and clearly indi-
cate the advantage of EMDeBruijn.

**Table 2:**
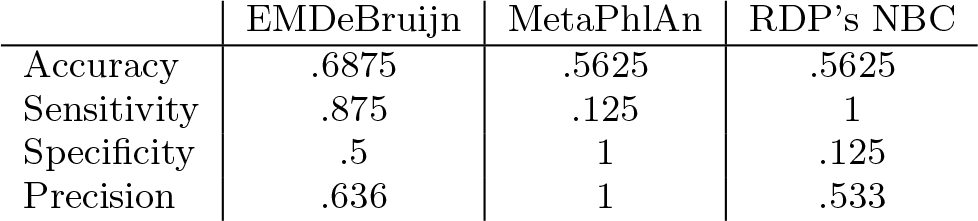
Binary classification results for the UPGMA trees derived from the two classification methods and EMDe-Bruijn.

The hierarchical clustering dendrograms associated to the three methods (mentioned in section 3.2 in the main text) are give in figure 13. See the associated table 1. Observe that in figure 13, EMDeBruijn partitions the samples into three clusters, each of which consists predominantly of case or control samples. Taking the three largest such clusters, we obtain the classification metrics contained in table 4. Note the significant improvement over table 1.

**Table 3:**
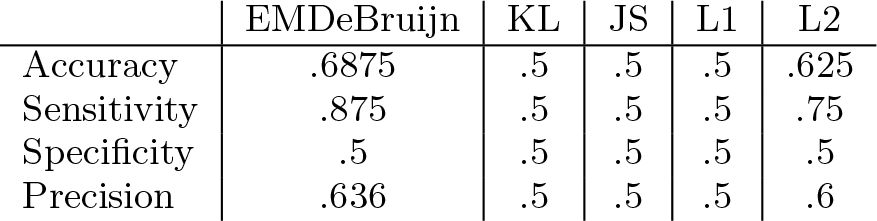
Binary classification results for the UPGMA trees derived from the five reference-free metrics. For *k*-mer size,*k* = 6 was used for EMDeBruijn, and *k* = 10 was used for the other methods. KL=Kullback-Leibler, JS=Jensen-Shannon.

**Table 4:** Classification results for the hierarchical clustering derived from the three classification methods. These measures result from taking the three largest clusters.

|  | EMDeBruijn | MetaPhlAn | RDP's NBC |
| --- | --- | --- | --- |
| Accuracy | .8125 | .5 | .562 |
| Sensitivity | .625 | .125 | .875 |
| Specificity | 1 | 1 | .25 |
| Precision | 1 | 1 | .538 |

**Figure 13:**
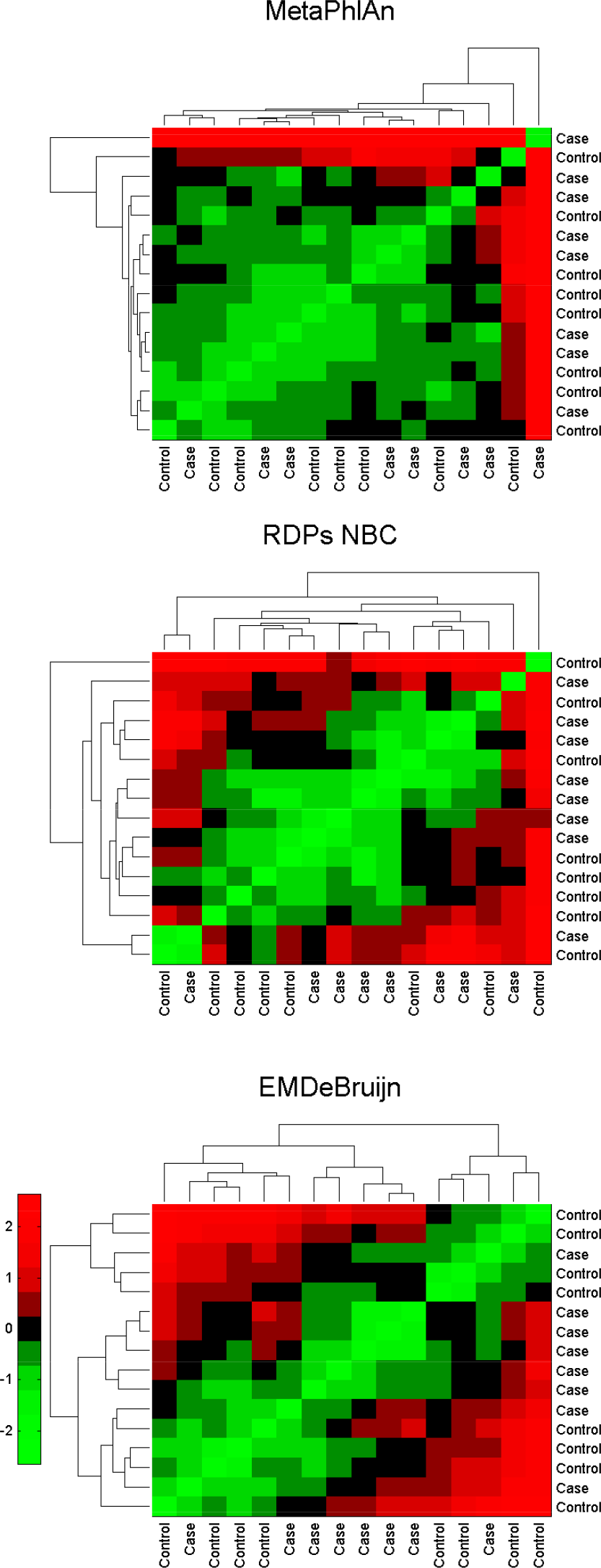
Dendrograms created via hierarchical clustering using the Jensen-Shannon divergence of the genus level classification for MetaPhlAn and RDP’s NBC, as well as the dendrogram created via hierarchical clustering based on the EMDeBruijn distance metric on the non-host reads. All distance matrices have been standardized.

**Figure 14:**
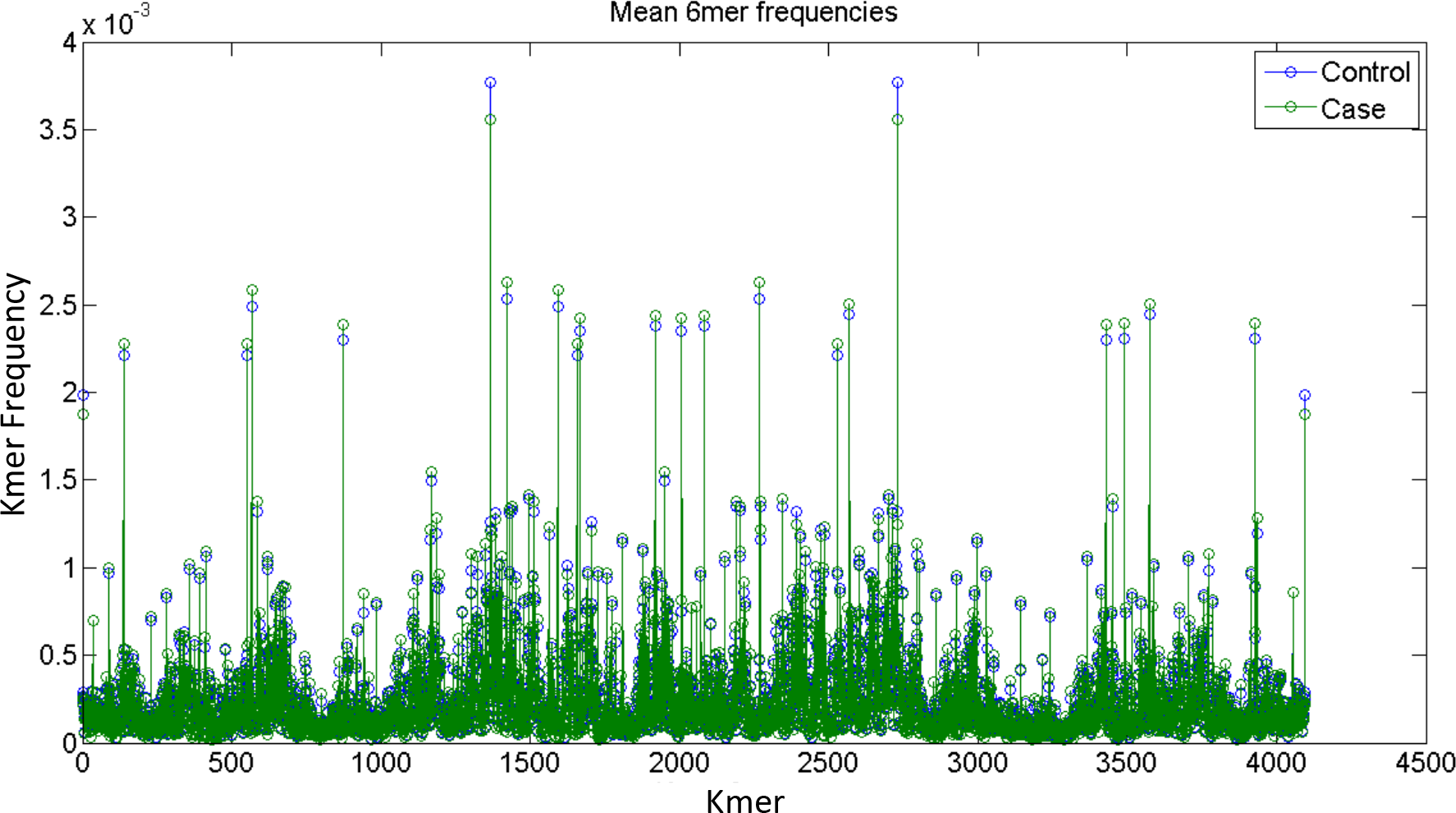
6-mers frequency plot.

